# MicroRNA-214 prevents pulmonary angiogenesis and alveolarization in neonatal rats with hyperoxia-mediated impairment of lung development by blocking PlGF-dependent STAT3 signaling pathway

**DOI:** 10.1101/2020.03.02.971572

**Authors:** Zhi-Qun Zhang, Xiao-Xia Li, Jing Li, Hui Hong, Xian-Mei Huang

## Abstract

In recent years, the roles of microRNAs (miRNAs) in pulmonary diseases have been widely studied and researched. However, the molecular mechanism by which miR-214 affects bronchopulmonary dysplasia (BPD) remains elusive and merits further exploration. Hence, this study aims to clarify the function of miR-214 in pulmonary angiogenesis and alveolarization in preterm infants with BPD. BPD neonatal rat model was induced by hyperoxia, and pulmonary epithelial cells were isolated from rats and exposed to hyperoxia. Gain- or loss-of-function experiments were performed in BPD neonatal rats and hyperoxic pulmonary epithelial cells. MiR-214 and PlGF expression in BPD neonatal rats, and eNOS, Bcl-2, c-myc, Survivin, α-SMA and E-cadherin expression in hyperoxic pulmonary epithelial cells were detected using RT-qPCR and western blot analysis. The interaction between PlGF and miR-214 was identified using dual luciferase reporter gene assay and RIP assay. ELISA was adopted to assess IL-1β, TNF-a, IL-6, ICAM-1 and Flt-1 expression in rats. Decreased miR-214 expression and elevated PlGF expression were evident in the lung tissues of neonatal rats with BPD. PlGF was a target of miR-214, and miR-214 downregulated PlGF to inactivate the STAT3 signaling pathway. miR-214 overexpression or PlGF silencing decreased apoptosis of hyperoxic pulmonary epithelial cells and declined pulmonary angiogenesis and alveolarization in BPD neonatal rats. Collectively, miR-214 can protects against pulmonary angiogenesis and alveolarization in preterm infants with BPD by suppressing PlGF and blocking STAT3 signaling pathway.

## INTRODUCTION

As a chronic lung disease, bronchopulmonary dysplasia (BPD) afflicts preterm infants and is characterized by retarded lung growth. It remains a principle cause of neonatal morbidity and triggers serious adverse consequences (13, 30). Around 45% of the preterm infants whose gestation was below 29 weeks are found to have BPD (25). It has already been documented that hyperoxia-induced acute lung injury is a key promoter of BPD pathogenesis in preterm infants (27). It is well known that lung alveolarization could cause incapacity of lungs to exchange gas effectively (23). However, the underlying molecular mechanisms are still largely unknown.

Through mediation of cellular proliferation, differentiation and metastasis, microRNAs (miRNAs or miRs) are involved in the pathogenesis of multiple diseases and thus have great potentials in serving as therapeutic targets (11). MiR-214, as a pivotal oncomiR, is upregulated in various kinds of diseases and cancers (32). In lung cancer, miR-214-3p has been demonstrated to downregulate fibroblast growth factor receptor 1 and to provide beneficial effects to patients (34). Also, downregulation of miR-214 could reverse the erlotinib resistance in non-small-cell lung cancer by upregulating its direct target gene LHX6 (15). In our study, placental growth factor (PlGF) was predicted to the target gene of miR-214. Moreover, it has been predicted in a prior study that PlGF might serve as a potential biomarker for BPD occurrence (36). Besides, PlGF could be increased under hyperoxic exposure and downregulating PlGF ameliorated the hyperoxia-induced lung impairment in neonatal rats (37). PlGF has also been proved to accelerate the phosphorylation of STAT3 (2). Moreover, neonatal exposure to hyperoxia has been found to lead to a significant increase of the signal transducer and activator of transcription 3 (STAT3) mRNA expression in pulmonary endothelial cells (5). In this regard, we hypothesized that a regulatory network of miR-214/PlGF/STAT3 signaling pathway may be involved in BPD. Therefore, the current study was conducted with the aim to verify the expected involvement of miR-214/PlGF/STAT3 axis in BPD, and to elucidate the underlying molecular mechanisms.

## MATERIALS AND METHODS

### Ethics statement

Animal experiment protocols were approved by the Experimental Animal Ethics Committee of Affiliated Hangzhou First People’s Hospital, Zhejiang University School of Medicine. All animal experiments were performed in accordance with the Guide for the Care and Use of Laboratory animals published by the US National Institutes of Health. Extensive efforts were made to ensure minimal suffering of animals during the study.

### Analysis of BPD gene expression dataset and bioinformatics prediction

The gene expression dataset GSE25293 of mouse BPD models was retrieved from the annotation platform GPL1261 in the Gene Expression Omnibus (GEO) database (https://www.ncbi.nlm.nih.gov/gds) and then analyzed using R language. The databases DIANA TOOLS (http://diana.imis.athena-innovation.gr/DianaTools/), miRWalk (energy < −30) (http://mirwalk.umm.uni-heidelberg.de), mirDIP (Integrated Score > 0.6) (http://ophid.utoronto.ca/mirDIP/), miRDB (http://www.mirdb.org) and miRSearch (https://www.exiqon.com/miRSearch) were used to analyze the intersected upstream miRNAs in human body. A box plot was drawn using R language to extract key miRNA expression data from the miRNA dataset GSE25293 from the annotation platform GPL11199 in the GEO database. Protein-protein interaction (PPI) analysis was performed on String website (https://string-db.org) to obtain proteins that could potentially bind to BPD. Cytoscape (https://cytoscape.org) was used to process the visual graphs of PPI analysis, and its downstream regulatory pathways were predicted based on existing literatures.

### Establishment of hyperoxia–induced BPD rat models

Twelve specific-pathogen-free (SPF) Sprague-Dawley (SD) rats with a gestational age of 15 days were purchased from Shanghai SLAC Laboratory Animal Co., Ltd. (Shanghai, China) and housed at 22 ± 3°C and with humidity of 60 ± 5% and circadian rhythm of 12 h. Each neonatal rat was housed individually and was self-delivered after 1 week of adaptive feeding. Then 60 neonatal rats within 12 h of birth difference after delivery were randomly grouped into the hyperoxia treatment group and the control group. And the hyperoxia-treated rats with BPD were then treated with miR-214 negative control (NC), miR-214 agomir, miR-214 NC + PlGF vector, and miR-214 agomir + PlGF (n = 12 in each treatment). After the recombinant adenoviruses were packaged, cloned, amplified, purified and titrated, the hyperoxia-induced BPD rats were subject to the following procedures as previously reported (3, 8): they were placed in an atmospheric oxygen box with polymethyl methacrylate. The oxygen was continuously inputted, and the fraction of inspired oxygen (Fio2) was maintained above 85%. Sodium lime was used to absorb CO_2_, and the temperature was set at 25 – 27°C with 50% – 70% humidity. The rats in the control group (Air group) were exposed to the air (with FiO2 of 21%), and the remaining experimental control conditions and operations were the same as those in the hyperoxia treatment group. The box was routinely opened for 30 min every day, and water and food were added and the litter was replaced. The mother rats were exchanged with the control group (to avoid the decreased feeding ability of mother rats because of oxygen toxicity). The rats in control group were placed in the same room, with the similar experimental control factors to those in the hyperoxia treatment group. Three newborn rats were randomly selected from the two groups by random number method at the 3rd, 7th and 14th days after the experiment began, and then received intraperitoneal injection of 90 mg/kg pentobarbital sodium for anesthesia. Next, the abdominal cavity was opened immediately, and the right lungs were taken out and placed in an RNase-free cryo vial (Eppendorf, Hamburg, Germany). After rapidly frozen with liquid nitrogen, the lungs were stored in a – 80°C refrigerator for subsequent reverse transcription quantitative polymerase chain reaction (RT-qPCR) and western blot analysis. Then 40 g/L paraformaldehyde was slowly injected to the rats through the left bronchus until the apex of lung was inflated, placed in an embedding box, and added with 40g/L paraformaldehyde solution for overnight fixation for subsequent detection.

### Enzyme-linked immunosorbent (ELISA) assay

The strips used in the experiment were equilibrated at room temperature for 20 min. Standard and sample wells were set separately, and 50 μL of standards (IL-1β, TNF-α and IL-6) at different concentration was added into the standard wells respectively. Then 10 μL of samples was added into the sample wells and 40 μL of sample dilution was then added to the samples. The blank wells were not subject to treatment. Next, 100 μL of horse radish peroxidase (HRP)-labeled antibody to be detected was added to the standard wells and the sample wells respectively, and the blank wells were not subject to treatment. The wells were sealed with microplate sealers, followed by incubation for 60 min in a 37°C incubator. After the liquid was discarded, the experimental strips were washed in full-automatic washing machine. Then 50 μL of substrate A and B were added to each well and incubated for 15 min a 37°C incubator in subdued light. Subsequently, 50 μL of the stop buffer was added to each well and allowed to stand for 15 min, after which the OD value of each well was measured at a wavelength of 450 nm.

### Hematoxylin-eosin (HE) staining

The lung tissues of rats in each group were fixed with 4% paraformaldehyde for 24 h, dehydrated with 80%, 90% and 100% ethanol and n-butanol respectively, immersed in a wax box at 60°C, dewaxed with xylene and hydrated. The sections were first stained with hematoxylin (Beijing Solarbio Science & Technology Co., Ltd., Beijing, China) for 2 min, then washed with distilled water for 1 min, stained with eosin for 1 min, rinsed with distilled water for 10 s, dehydrated with gradient ethanol, cleared with xylene, and fixed with neutral rubber. Finally, the morphological changes of lung tissues were observed and analyzed under an optical microscope (XP-330, Bingyu Optical Instrument Co., Ltd., Shanghai, China).

### Alveolar epithelial cell isolation using Immunomagnetic bead

The fetal rats were taken out by cesarean section from the SD rats with a gestational age of 15 days under sterile condition and were transected at the chest, after which the lungs were taken out and put in pre-cooled phosphate buffer solution (PBS) to remove residual non-lung tissues. Digestion, filtration, centrifugation, sorting and so on were carried out according to the experimental operations. CD14 magnetic beads were added to the cells (20 μL/10^7^ cells), mixed well, and incubated at 4 – 8°C for 15 min. The cells were washed with buffer solution (1 mL/10^7^ cells) and centrifuged at 1500 r/min for 10 min, with the supernatant completely removed. The cells were resuspended in 500 μL of buffer solution, and the cell suspension was added to a MS separation column. The unlabeled cells that had flowed out first were collected, which were negative cells. The MS separation column was washed with 1500 μL of buffer solution. The separation column was removed from the magnetic field and the cells retained on the column were quickly eluted with 1 mL of buffer solution. These cells were magnetically labeled positive cells. Under a modified Barthel microscope, more dark particles in the cytoplasm were visible with characteristic eosinophils observed, which presented with obvious microvilli in lamellar bodies and cell membranes. This indicated that type II alveolar epithelial cells in the fetal rats were successfully isolated.

### Development of hyperoxia-provoked cell injury models

Pulmonary epithelial cells after 2 days of growth were exposed to air (control) or hyperoxia. The cells exposed to air and hyperoxia were placed in a closed oxygen chamber with 21% oxygen volume fraction. The hyperoxic cells were further transfected with plasmids of miR-214 NC, miR-214 mimic, miR-214 NC + PlGF vector, miR-214 NC + PlGF, and miR-214 mimic + PlGF. Then the recombinant adenoviruses were packaged, cloned, amplified, purified and titrated. The pulmonary epithelial cells were infected with these adonoviruses and placed in a closed oxygen chamber with 85% oxygen volume fraction.

### Transmission electron microscope

After the pulmonary epithelial cells were centrifuged at 10,000 rpm for 10 min, the supernatant was discarded and the cells were fixed in 4% glutaraldehyde at 4°C for more than 2 h. The cells were then washed with 0.01 M PBS three times, fixed with 1% osmium tetroxide for 2 h, and dehydrated with gradient ethanol and acetone. The cells were then immersed with epoxy resin, embedded and polymerized, and then made into semi-thin sections with a thickness of 0.5 μm. The sections were positioned under a microscope, stained with uranyl acetate and lead citrate, and observed and photographed under a transmission electron microscope (H-7500).

### RNA binding protein immunoprecipitation (RIP)

The pulmonary epithelial cells were washed twice with cold PBS, then added with 10 mL of PBS, scraped off with a cell scraper and transferred into a centrifuge tube. The cells were centrifuged at 1500 rpm for 5 min at 4°C, then added with RIP Lysis Buffer, mechanically dissociated into and mixed thoroughly, and lysed on ice for 5 min to prepare cell lysate. Next, 50 μL of magnetic beads were added into each tube and mixed well, after which 0.5 mL RIP Wash Buffer was added to rinse the magnetic beads, and 100 μL RIP Wash Buffer was added to to resuspend beads. Then 5 μg of Ago2 antibody was added to the tubes and incubated in rotation for 30 min at room temperature. The supernatant was discarded and the beads were washed twice with 0.5 mL RIP Wash Buffer for subsequent experiments. 900 μL of RIP Immunoprecipitation Buffer was added to the magnetic bead-antibody mixture, after which the centrifugation was carried out at 14000 rpm for 10 min at 4°C. The supernatant was collected and transferred into a new eppendorf (EP) tube, and then 100 μL of the supernatant was taken into the tube containing the magnetic bead-antibody. 1.0 mL served as the final volume of the immunoprecipitation reaction, and incubation was conducted overnight at 4°C. Then the magnetic beads were washed 6 times with 0.5 mL RIP Wash Buffer, 150 μL of proteinase K buffer was added, and the RNA was purified by 30 min of incubation at 55°C. The RNA was extracted by a conventional TRIzol method followed by RT-qPCR detection.

### Giemsa staining

The rats were anesthetized by intraperitoneal injection of 1% sodium pentobarbital solution, and fixed on a simple operation table. The thoracic cavities of rats were exposed after being euthanized by exsanguination, after which the pleural tissues around the trachea were bluntly separated, the trachea was fully exposed and a needle was used to stab at the 1/3 of the trachea. The tip of the 18G indwelling needle was previously trimmed into a hernia type, and the tracheal intubation was performed along the trial point and ligated with a surgical line. 1 mL of pre-cooled sterile normal saline was used to perfuse the lung tissues of the rats for three times. The bronchoalveolar lavage fluid (BALF) was collected and stored in a pre-cooled EP tube and subsequently centrifuged at 1200 r/min for 20 min at 4°C. The supernatant was stored in a −80°C freezer for subsequent use. The cell precipitate was resuspended in 100 μL of PBS, smeared with 50 μL of the suspension, and stained with Swiss Giemsa. The number of neutrophils was counted under an oil microscope and the total number of cells in 10 μL of the suspension was counted by a hemocytometer.

### Immunohistochemistry

The specimen was fixed in 10% formaldehyde, and embedded into paraffin and continuously made into 4 μm sections. Then the tissue sections were placed in a 60°C oven for 1 h, dewaxed by xylene in conventional manner, then dehydrated with gradient alcohol, and incubated in 3% H_2_O_2_ (Sigma-Aldrich, Shanghai, China) for 30 min at 37°C. Then the tissue sections were boiled in 0.1 M citrate buffer solution for 20 min at 95°C after PBS rinse, cooled to room temperature, and rinsed with PBS again. The sections were blocked with 10% normal goat serum for 10 min at 37°C followed by rabbit anti-eNOS (AF0096, Affinity) incubation at 4°C for 12 h. After PBS wash, the sections were treated with biotin-labeled goat anti-rabbit secondary antibody at room temperature for 10 min. After thoroughly washed, S-A/HRP was added to react at room temperature for 10 min. The tissue sections were developed using diaminobenzidine (DAB) away from light at room temperature for 8 min. Then the tissues were rinsed with tap water, counter-stained with hematoxylin, dehydrated, cleared, blocked, and observed under a light microscope. The number of positive cells was counted using image analysis software (Nikon Corporation, Tokyo, Japan). Five fields of equal area were selected from each section, and the proportion of positive cells was calculated with the average value calculated. The cells with apparent brown or brownish yellow particles in the cytoplasm were positive cells. The experiment was repeated three times.

### Dual luciferase reporter gene assay

The target gene of miR-214 was analyzed on biological prediction website, and dual luciferase reporter gene assay was performed to further verify whether PlGF was a direct target of miR-214. In short, the artificially synthesized PlGF 3’ untranslated regions (UTR) gene fragment was constructed into pMIR-reporter (Promega, Madison, WI, USA). A complementary sequence with mutation of the seed sequence was designed based on the wild type (WT) of PlGF and constructed into the pMIR-reporter reporter plasmid. The correctly sequenced luciferase reporter plasmids WT and MUT were respectively co-transfected with miR-214 mimic and miR-214 NC into HEK293T cells. After 48 h of transfection, cells were collected and lysed, and the luciferase activity was measured using Dual-Luciferase Reporter Assay System (Promega, Madison, WI, USA). The experiment was repeated three times.

### RT-qPCR assay

Total RNA was extracted from cells using the Trizol kit (Invitrogen Inc., Carlsbad, CA, USA) and reverse transcribed into cDNA according to the instructions of TaqMan MicroRNA Assays Reverse Transcription Primer (4427975, Applied Bio-systems, Foster City, CA,USA). The reverse transcribed cDNA was diluted to 50 ng/μL. The expression of relevant genes was analyzed. U6 gene was taken as an internal reference of miRNA, and glyceraldehyde-3-phosphate dehydrogenase (GAPDH) was taken as an internal reference of other genes. The fold changes were calculated using relative quantification (the 2^-ΔΔCt^ method). The primers used are shown in Table 1.

**Table 1.**
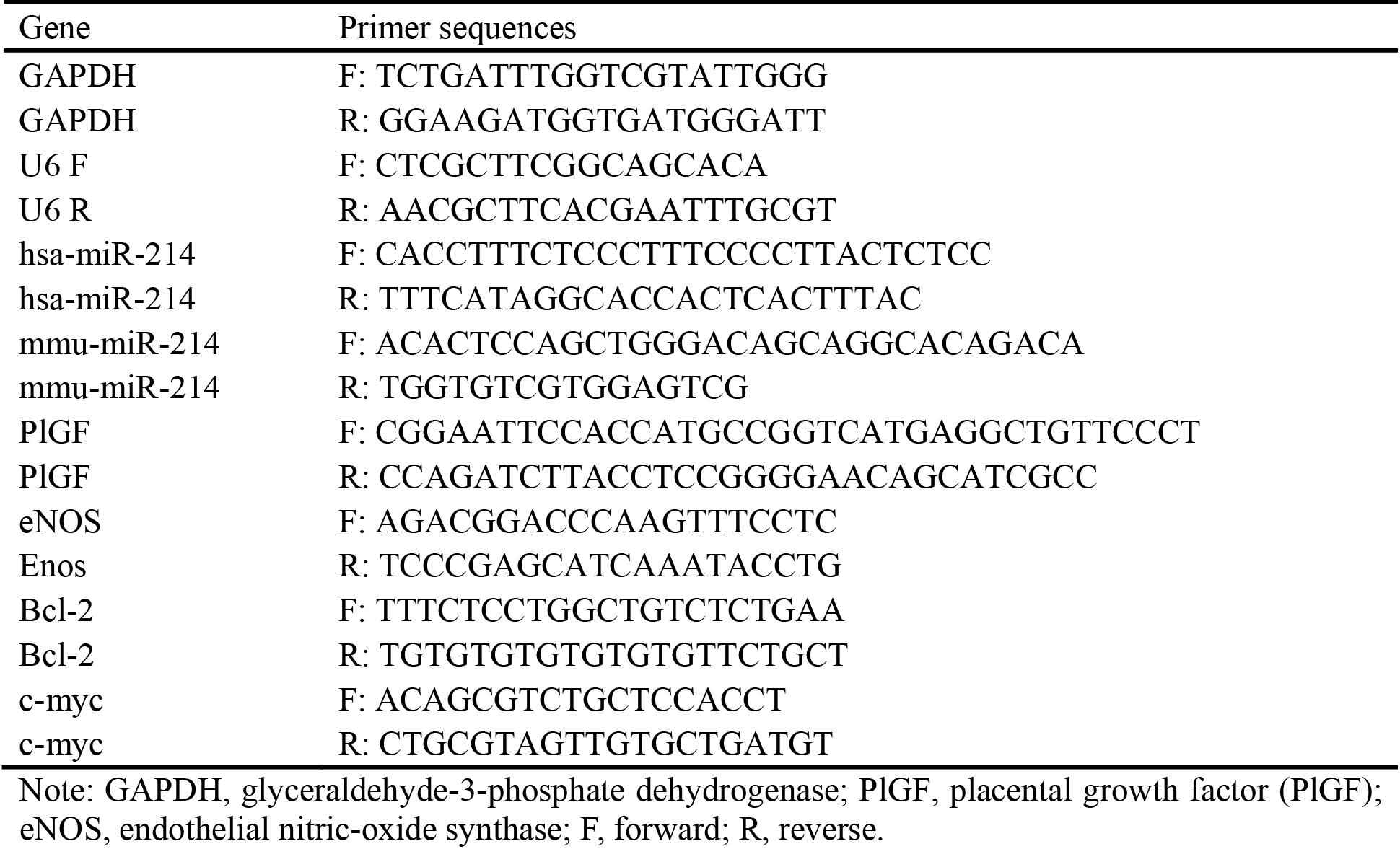
Primer sequences of RT-qPCR

### Western blot assay

The lung tissues or cells were washed twice with PBS, added with lysis buffer, shaken on a vortex agitator, and centrifuged at 12000 r/min for 30 min at 4°C to remove tissues or cell debris. The supernatant was taken and the total protein concentration was measured using a bicinchoninic acid (BCA) kit. 50 μg of protein was subjected to 10% sodium dodecyl sulfate polyacrylamide gel electropheresis and transferred to polyvinylidene fluoride membranes by wet transfer method. After blocked with 5% skim milk powder at room temperature for 1 h, the PVDF membrane was then incubated with diluted primary antibodies Survivin (# 2808S, CST, Danvers, MA, USA), GAPDH (#5174, CST, Danvers, MA, USA), B-cell lymphoma-2 (Bcl-2, #3498, CST, Danvers, MA, USA), c-myc (#13987, CST, Danvers, MA, USA), PlGF (ab74778, Abcam, Cambridge, UK), E-cadherin (ab11512, Abcam, Cambridge, UK) and α-smooth muscle actin (α-SMA, ab32575, Abcam, Cambridge, UK), and then diluted according to the instructions. The membrane was washed 3 times with TBST and then incubated with HRP-labeled secondary antibody for 1 h. After rinsed with TBST, the membrane was placed on a clean glass plate. The immunocomplexes on the membrane were visualized using enhanced chemiluminescence (ECL) fluorescence detection kit (BB-3501, Amersham, Little Chalfont, UK), and band intensities were quantified using a Bio-Rad image analysis system and Quantity One v4.6.2 software. The ratio of the gray value of the target band to GAPDH was representative of the relative protein expression.

### Statistical analysis

Data analyses were conducted using SPSS 21.0 (IBM Corp, Armonk, NY, USA). Measurement data were described using mean ± standard deviation. An unpaired t-test was conducted to compare the data obeying normal distribution and homogeneity of variance between two groups. Data comparisons between multiple groups were performed using one-way analysis of variance (ANOVA), followed by a Tukey’s multiple comparisons posttest. Data comparisons at different time points were performed by repeated measures ANOVA, followed by a Bonferroni post hoc test for multiple comparisons. Pearson correlation was used to analyze the relationship between two indicators. A value of *p* < 0.05 was considered to be statistically significant.

## RESULTS

### miRNA and mRNA expression profiles in BPD

There has been literature showing that PlGF is an important gene participating in BPD in preterm infants, but the regulatory pathway of this gene is still unknown and possesses great research potential (14). A boxplot shown in Figure 1A illustrating that PlGF was highly expressed in BPD model. The predicted results from DIANA TOOLS, miRWalk (energy < −30), mirDIP (Integrated Score > 0.6), miRDB and miRSearch databases revealed that the upstream miRNA number of PlGF (actually PGF was used during prediction) were 80, 196, 5, 58 and 39 respectively. Only one intersected miRNA was obtained shown by the Venn map, which was miR-214 (Fig. 1B). Besides, the binding site suggested by DIANA Tools also indicated the potential interaction between PlGF and miR-214-3p (Fig. 1C). The boxplot on miR-214 expression drawn using R language showed that miR-214 was downregulated in BPD neonatal rats induced by hyperoxia (Fig. 1D). Existing studies have shown that miR-214-3p is a miRNA related to lung cancer and is downregulated in lung tumor tissues (6). PPI analysis indicated that PlGF (PGF) may be related with several signaling pathways (Fig. 1E), and some literature has shown that high expression of PlGF can promote the expression of STAT3 (20, 22). Therefore, we hypothesized that miR-214 can regulate the expression of PlGF and further regulate the STAT3 signaling pathway, thus affecting the progression of BPD in preterm infants.

**Fig. 1.**
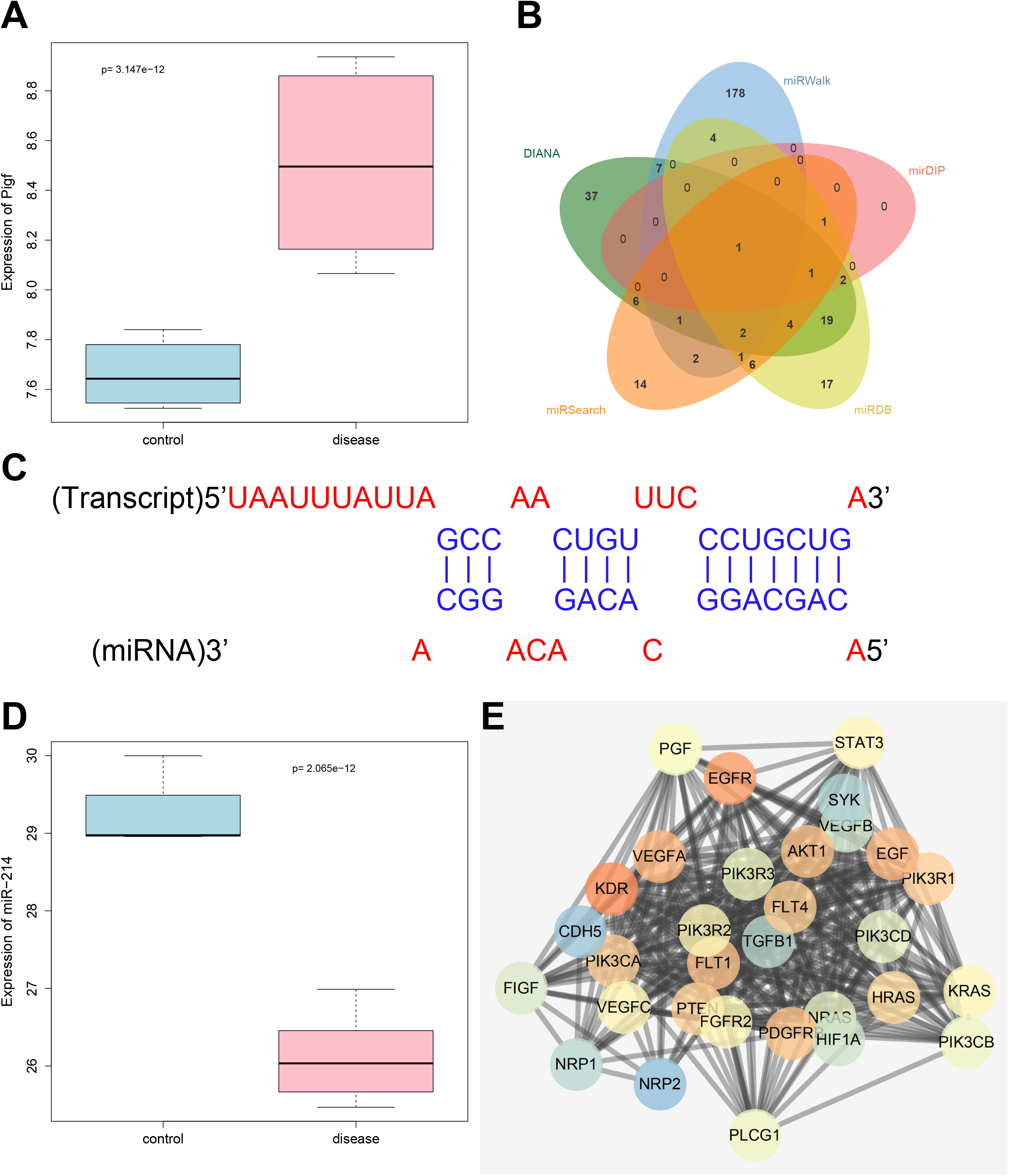
PlGF/miR-214/STAT3 axis is predicted to be involved in BPD. *A*, PlGF expression in the dataset GSE25293-GPL1261, in which the blue box on the left indicates the expression of normal samples, and the red box on the right indicates the expression of BPD samples. *B*, The upstream miRNAs of PlGF predicted by DIANA Tools, miRWalk, mirDIP, miRDB and miRSearch databases. *C*, The binding site of PlGF and miR-214 predicted by DIANA Tools. *D*, miR-214 expression in the dataset GSE25293-GPL11199, in which the blue box on the left indicates the expression of normal samples, and the red box on the right indicates the expression of BPD samples. *E*, PPI analysis of PlGF (PGF), in which the redder gene sphere indicates greater importance, and the bluer gene sphere indicates a lower importance.

### miR-214 is downregulated in the lung tissues of hyperoxia-induced BPD neonatal rats

In order to verify miR-214 expression in the lung tissues of neonatal rats exposed to hyperoxia, neonatal rats were selected for experiments. After BPD models were established, miR-214 expression in the lung tissue of neonatal rats exposed to hyperoxia was first detected by RT-qPCR. The results showed that on 3rd, 7th, and 14th days, miR-214 expression in lung tissue of neonatal rats with BPD was lower than that of normal neonatal rats (*p* < 0.05; Fig. 2A). PlGF expression was detected by RT-qPCR and western blot analysis, which showed that PlGF expression in the lung tissue of neonatal rats with BPD were higher than that in normal neonatal rats on 3rd, 7th, and 14th days (*p* < 0.05; Fig. 2B-C). Correlation analysis showed a negative correlation between miR-214 and PlGF expressed in lung tissues (*p* < 0.05; Fig. 2D). The above results suggested that expression level of miR-214 was decreased and PlGF was elevated in lung tissue of neonatal rats exposed to hyperoxia, and there was a certain correlation between them.

**Fig. 2.**
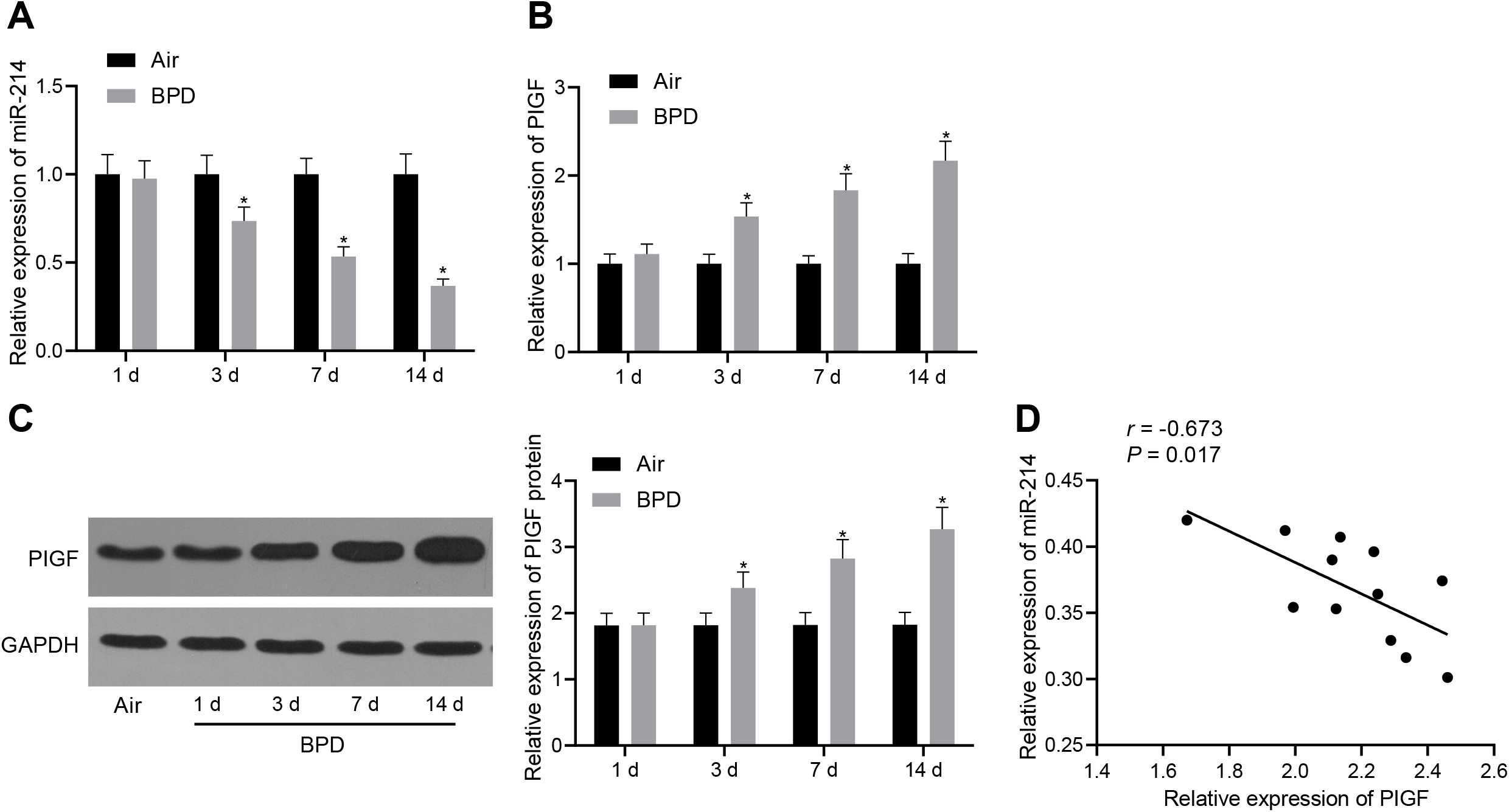
Decreased miR-214 expression and increased PlGF expression are found in the lung tissues of neonatal rats with BPD. *A*, miR-214 expression in neonatal rats with BPD detected by RT-qPCR normalized to U6. *B*, mRNA level of PlGF in neonatal rats with BPD detected by RT-qPCR normalized to GAPDH. *C*, Western blot analysis of PlGF protein expression in neonatal rats with BPD normalized to GAPDH. *D*, Correlation analysis of miR-214 and PlGF expression. The data were measurement data and expressed as mean ± standard deviation. * *p* < 0.05 *vs.* neonatal rats exposed to air. Data comparisons at different time points were performed by repeated measures ANOVA, followed by a Bonferroni post hoc test for multiple comparisons. n = 12.

### Overexpression of miR-214 blocks pulmonary angiogenesis and alveolarization in neonatal rats with BPD in vivo

To validate the effect of miR-214 overexpression on neonatal rats with BPD, we established hyperoxia-induced neonatal rats with BPD and we injected them with miR-214 NC and miR-214 agomir. Then, ELISA, immunohistochemistry, Giemsa staining, and HE staining were performed respectively. When compared with neonatal rats exposed to air, the BPD neonatal rats exhibited increased levels of inflammatory factors (IL-1β, TNF-α, IL-6, Flt-1 and ICAM-1) (*p* < 0.05; Fig. 3A-D), elevated eNOS expression (Fig. 3E), and increased number of macrophages (Fig. 3F) (all *p* < 0.05), which was reversed by miR-214 agomir treatment (*p* < 0.05). HE staining revealed that compared with neonatal rats exposed to air, BPD neonatal rats showed reduced alveoli number and simplified structure, the alveolar wall ruptured and merged into pulmonary bullae, the pulmonary microvessel density was decreased, and the ratio of alveolar area/pulmonary septal area was increased (*p* < 0.05). While after miR-214 treatment, the alveolar structure of BPD neonatal rats exhibited a contrasting trend in the aforementioned factors (*p* < 0.05; Fig. 3G-H). The above results indicated that overexpression of miR-214 can prevent pulmonary angiogenesis and alveolarization in neonatal rats with BPD.

**Fig. 3.**
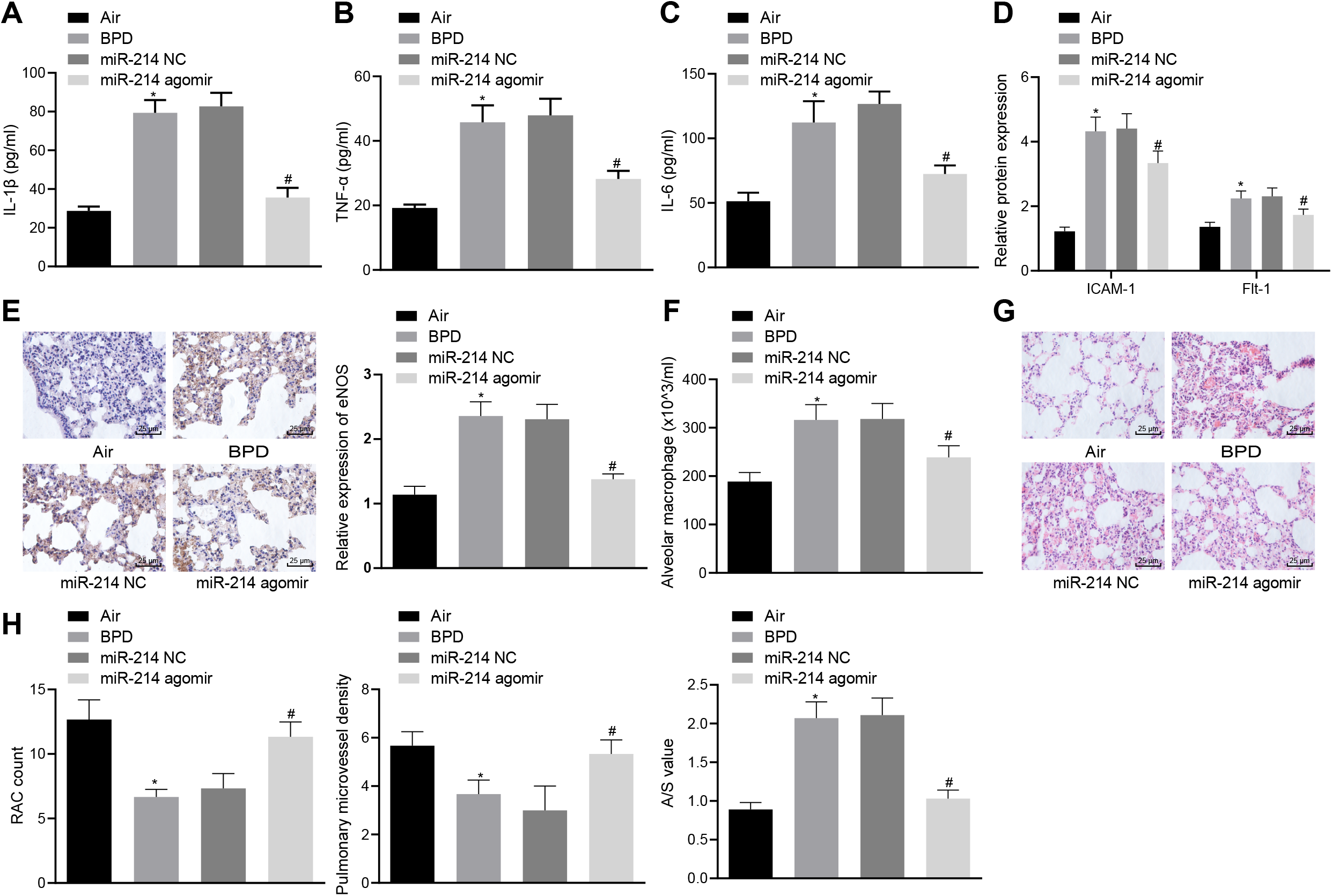
Overexpressed miR-214 represses pulmonary angiogenesis and alveolarization in neonatal rats with BPD. BPD neonatal rats were injected with miR-214 NC and miR-214 agomir. *A-C*, ELISA detection of the expression of inflammatory factors IL-1β, TNF-α and IL-6 in rats. *D*, Detection of ICAM-1 and Flt-1 levels in rats by ELISA. *E*, Immunohistochemistry of eNOS expression in pulmonary microvascular endothelium of rats (× 400). *F*, Giemsa staining of the number of macrophages in rats. *G*, HE staining of pulmonary microvascular (× 400). *H*, HE staining showing the number of alveoli, pulmonary microvessel count, and the growth of alveolar. The data were measurement data and expressed as mean ± standard deviation. * *p* < 0.05 *vs.* neonatal rats exposed to air. #*p* < 0.05 *vs*. BPD neonatal rats treated with miR-214 NC. Data comparisons among multiple groups were performed using one-way ANOVA and Tukey’s for post hoc test, and n = 12.

### Overexpression of miR-214 decreased the pulmonary epithelial cell apoptosis in vitro

To elucidate the effect of miR-214 on pulmonary epithelial cells, pulmonary epithelial cells were obtained from rats, and their ultrastructures after transfection were observed under a transmission electron microscope. The results showed that compared with pulmonary epithelial cells exposed to air, the pulmonary epithelial cells treated with hyperoxia displayed destroyed cytoplasmic lamellar structure of the alveolar epithelium, with relatively large vacuoles formed and gap between blood-air barriers increased, which can be rescued by miR-214 mimic transfection (Fig. 4A). Next, Western blot analysis was performed to assess the expression of apoptosis-related genes (Survivin, Bcl-2 and c-myc). As depicted in Figure 4B-C, in hyperoxic pulmonary epithelial cells, Survivin, Bcl-2 expression was decreased and c-myc expression was increased, while miR-214 mimic treatment triggered an opposite trend of the gene expression level. Also, α-SMA and E-cadherin expression in pulmonary epithelial cells was detected. α-SMA expression was increased while E-cadherin expression was decreased in hyperoxic pulmonary epithelial cells. While in the miR-214-treated cells, we observed decreased α-SMA expression and elevated E-cadherin expression (Fig. 4D-E). Taken together, overexpression of miR-214 can attenuate pulmonary epithelial cell alteration in BPD rats.

**Fig. 4.**
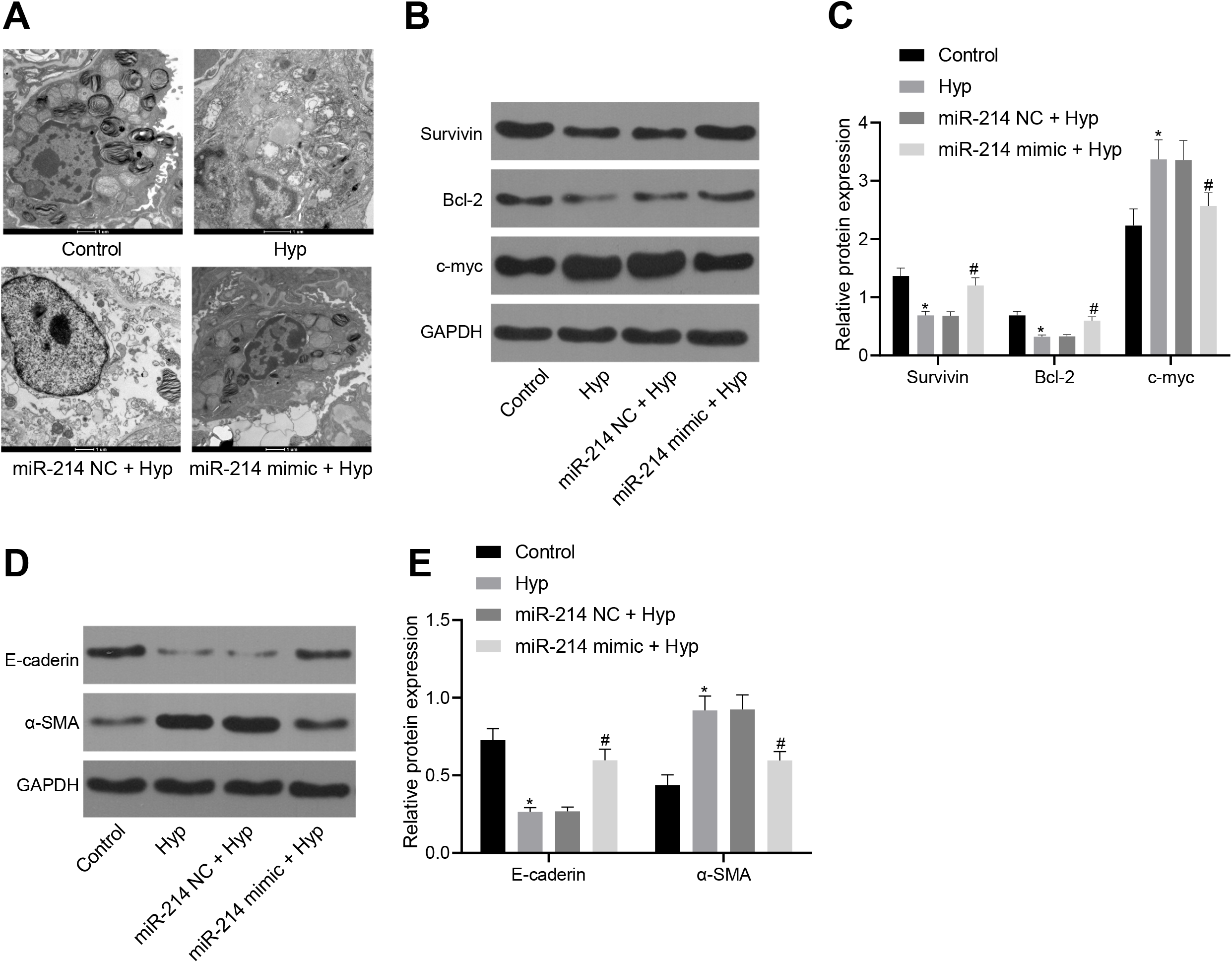
miR-214 overexpression exerts positive effects on the embryonic pulmonary epithelial cells of rats with BPD. Hyperoxic pulmonary epithelial cells were transfected with miR-214 NC or miR-214 mimic. *A*, The ultrastructure of alveolar epithelial cells under a transmission electron microscope (× 10000). *B-C*, Western blot analysis of Survivin, Bcl-2 and c-myc proteins in embryonic pulmonary epithelial cells. *D-E,* Western blot analysis of E-cadherin and α-SMA proteins in embryonic pulmonary epithelial cells. The data were measurement data and expressed as mean ± standard deviation. * *p* < 0.05 *vs.* pulmonary epithelial cells exposed to air. # *p* < 0.05 vs. hyperoxic pulmonary epithelial cells treated with miR-214 NC. Data comparisons were performed by one-way ANOVA, followed by a Tukey’s post hoc test for multiple comparisons and the experiment was repeated three times.

### PlGF is a target gene of miR-214

Then, the regulatory mechanism of miR-214 was explored. Bioinformatics analysis using an online prediction software revealed a complementary sequence of PlGF within the sequence of miR-214 (Fig. 5A). Dual luciferase reporter gene assay verified that co-transfection of miR-214 mimic with the 3’-UTR of WT-PlGF showed reduced luciferase activity, whereas co-transfection of miR-214 mimic with the 3’-UTR of MUT-PlGF showed no significant difference (Fig. 5B). RT-qPCR and western blot analysis presented that the mRNA and protein level of PlGF were reduced in cells treated with miR-214 mimic (Fig. 5C-D). RIP experiment further confirmed that the enrichment of miR-214 and PlGF was higher in Ago2 group than that in IgG (Fig. 5E). Collectively, miR-214 targeted and regulated the expression of PlGF.

**Fig. 5.**
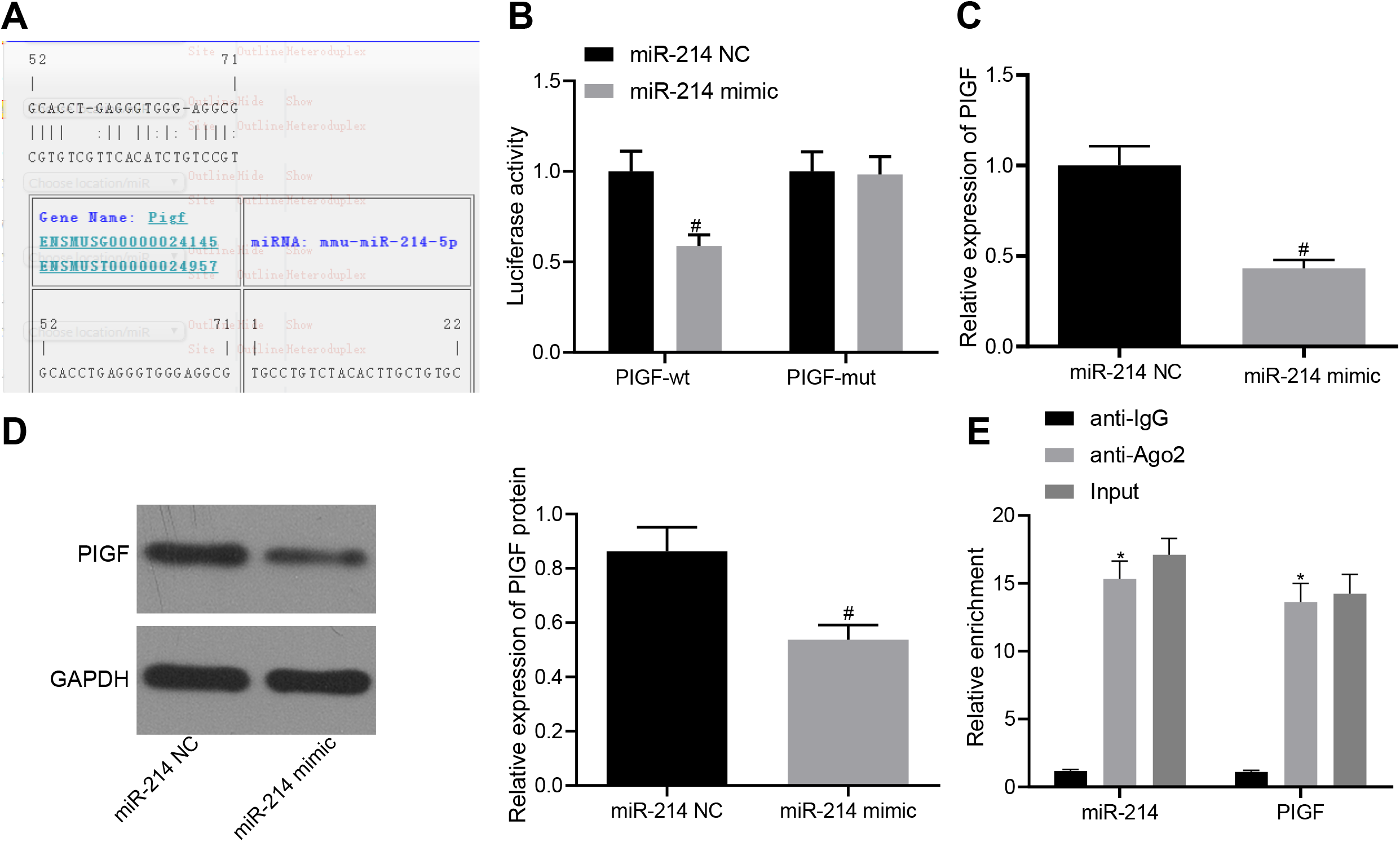
PlGF is identified to be a target of miR-214. *A*, The predicted binding sites of miR-214 and PlGF. *B*, Dual luciferase reporter gene assay analysis of the binding of miR-214 to PlGF. *C*, RT-qPCR assay detection of mRNA level of PlGF after miR-214 overexpression. *D*, Western blot analysis of PlGF after miR-214 overexpression. *E*, RIP detection of the binding percentage of miR-214 and PlGF to Ago2 normalized to IgG binding. The data were measurement data and expressed as mean ± standard deviation. * *p* < 0.05 *vs.* anti-IgG. # *p* < 0.05 vs. cells transfected with miR-214 NC. An independent sample t-test was used for comparison between two groups. Data comparisons among multiple groups were performed by one-way ANOVA, followed by a Tukey’s post hoc test for multiple comparisons. The experiment was repeated three times.

### Overexpressed miR-214 disrupts pulmonary angiogenesis and alveolarization by inactivating PlGF-dependent STAT3 signaling pathway in neonatal rats with BPD

The downstream regulatory pathway of PlGF was predicted using KEGG and GeneMANIA databases and the results showed that PlGF was closely related to the STAT3 signaling pathway (Fig. 6A). To examine the regulatory role of STAT3 signaling pathway in BPD, we used western blot analysis to detect the activation of the STAT3 signaling pathway in lung tissue of neonatal rats exposed to hyperoxia. The results showed that STAT3 phosphorylation was increased in the lung tissue of neonatal rats exposed in hyperoxia, which suggested that STAT3 signaling pathway was activated hyperoxia treated neonatal rats (Fig. 6B). The neonatal rats with BPD were then respectively treated with miR-214 NC + PlGF vector, miR-214 NC + PlGF, or miR-214 agomir + PlGF. Then, western blot analysis, ELISA, immunohistochemistry, giemsa staining and HE staining were performed. The results showed that compared with BPD rats treated with miR-214 NC + PlGF vector, the BPD rats treated with miR-214 NC + PlGF presented with increased expression of phosphorylated STAT3/STAT3 (Fig. 6C), levels of IL-1β, TNF-α, IL-6, ICAM-1 and Flt-1 (Fig. 6D-E), eNOS expression (Fig. 6F) and number of macrophages (Fig. 6G), which was abrogated by treatment of miR-214 agomir + PlGF (all *p* < 0.05). Moreover, compared with BPD rats co-treated with miR-214 NC and PlGF vector, BPD rats treated with miR-214 NC + PlGF displayed reduced number of alveoli and simplified structure. The alveolar wall ruptured to form a large pulmonary vesicle with declined pulmonary microvessels density and elevated ratio of alveolar area/pulmonary septal area (A/S) (*p* < 0.05; Fig. 6H-I), and we observed a opposite trend in alveolar alterations in BPD rats treated with miR-214 agomir + PlGF. These results suggested that miR-214 targeted PlGF to inhibit the STAT3 signaling pathway, thus blocking pulmonary angiogenesis and alveolarization in neonatal with BPD.

**Fig. 6.**
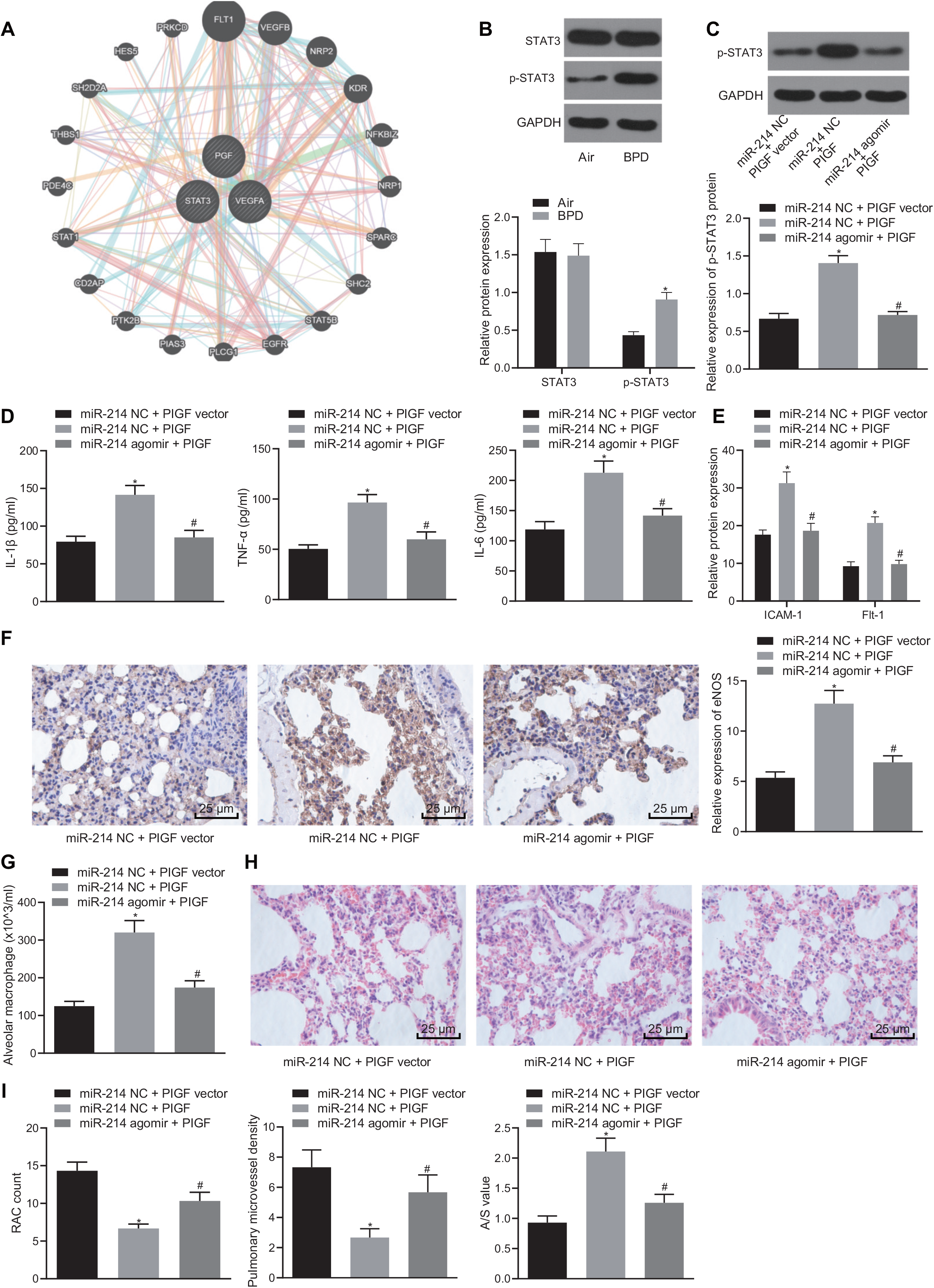
Overexpressed miR-214 blocked the effect of activated STAT3 signaling pathway on pulmonary angiogenesis and alveolarization by inhibiting PlGF in neonatal rats with BPD. *A*, The KEGG and GeneMANIA database were used to predict the downstream regulatory pathways of PlGF. *B*, Western blot analysis of the activation of STAT3 signaling pathway in BPD neonatal rats normalized to GAPDH. The neonatal rats with BPD were then respectively treated with miR-214 NC + PlGF vector, miR-214 NC + PlGF, or miR-214 agomir + PlGF. *C*, Western blot analysis of the expression of phosphorylated STAT3 in BPD neonatal rats normalized to GAPDH. *D*, Detection of the levels of inflammatory factors IL-1β, TNF-α and IL-6 by ELISA. *E*, Detection of levels of ICAM-1 and Flt-1 by ELISA. *F*, Immunohistochemistry analysis of eNOS expression (× 400). *G*, Giemsa staining detection of the number of macrophages. *H*, HE staining of the formation of pulmonary microvascular (× 400). *I*, HE staining of the number of alveoli and pulmonary microvessel, and alveolar growth. The data were measurement data and expressed as mean ± standard deviation. * *p* < 0.05 *vs.* BPD rats treated with miR-214 NC + PlGF NC. # *p* < 0.05 vs. BPD rats treated with miR-214 NC + PlGF. An independent sample *t*-test was used for comparison between two groups. Data comparisons among multiple groups were performed by one-way ANOVA, followed by a Tukey’s post hoc test for multiple comparisons. n = 12.

### Overexpressed miR-214 reduces pulmonary epithelial cell apoptosis via inactivation of PlGF-dependent STAT3 signalingpathway.in vitro

To study the effect of miR-214 on pulmonary epithelial cells of rats with BPD, pulmonary epithelial cells were obtained from rats and their ultrastructures after transfection were observed under a transmission electron microscope. The result showed that, compared with the cells co-transfected with miR-214 NC and PlGF vector, in the rats following transfection with miR-214 NC + PlGF, the cytoplasmic lamellar structure of the alveolar epithelium was destroyed, large vacuoles were formed, the blood-air barriers gap was increased (Fig. 7A). Moreover, there was a reduction in the expression of anti-apoptotic proteins Survivin and Bcl-2 (Fig. 7B-C) as well as α-SMA (Fig. 7D-E), while an increase in the expression of pro-apoptotic protein c-myc (Fig. 7B-C) and E-cadherin (*p* < 0.05; Fig. 7D-E). When compared with the rats co-transfected with miR-214 NC and PlGF, in the rats co-transfected with miR-214 mimic and PlGF, the cytoplasmic lamellar structure of the alveolar epithelium and the gap of the blood-air barrier were improved (Fig. 7A), Survivin and Bcl-2 expression were significantly increased (Fig. 7B-C), and α-SMA expression was significantly elevated (Fig. 7D-E), together with reduced expression of c-myc (Fig. 7B-C) and E-cadherin (*p* < 0.05; Fig. 7D-E). These results indicated that overexpression of miR-214 targeting PlGF can inhibit the effect of STAT3 signaling pathway on rat bronchial embryonic pulmonary epithelial cells.

**Fig. 7.**
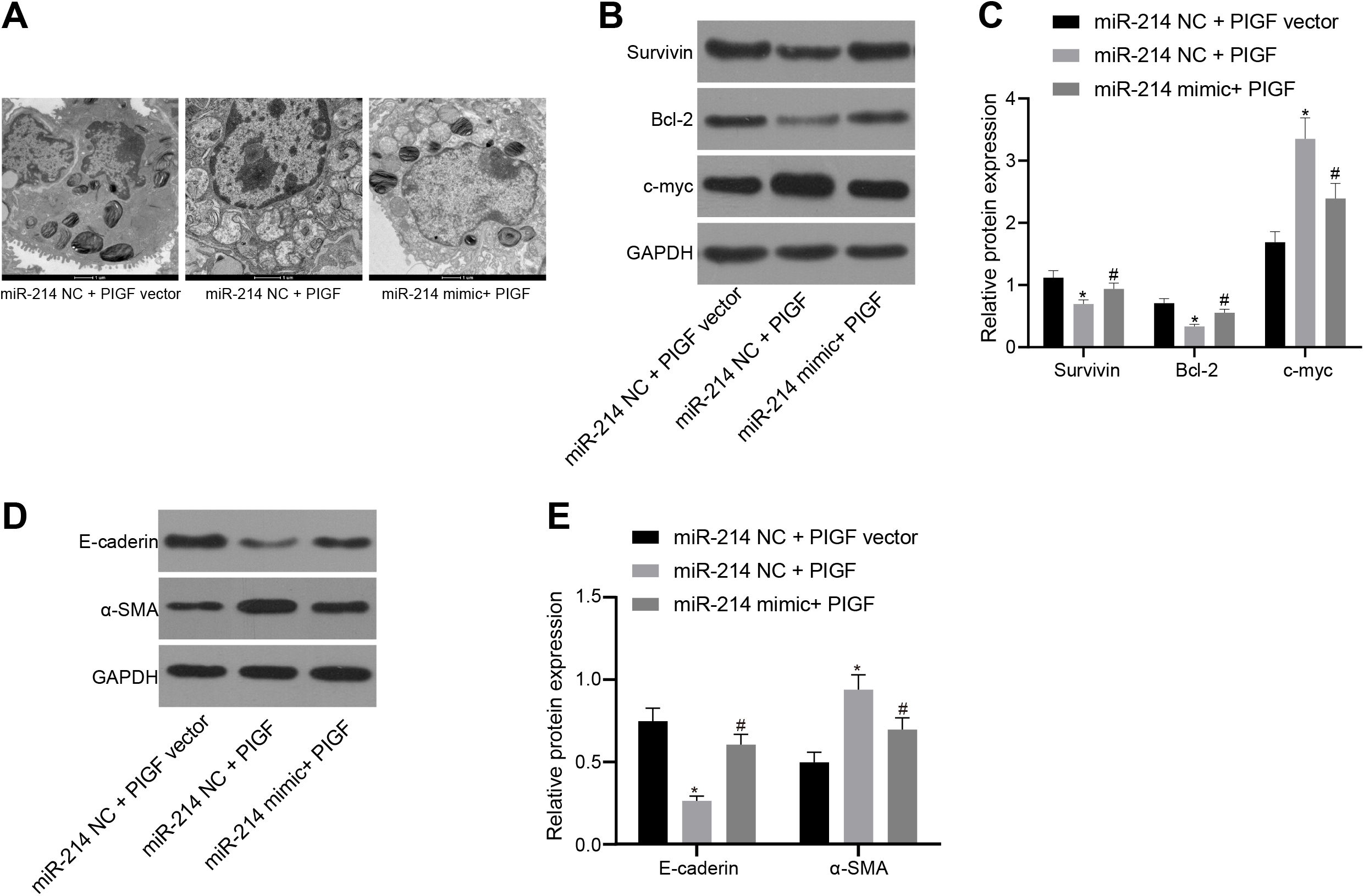
Overexpressed miR-214 blocked the effect of activated STAT3 signaling pathway on the bronchial embryonic pulmonary epithelial cells by inhibiting PlGF. *A*, The ultrastructure of alveolar epithelial cells under a transmission electron microscope (× 10000). *B-C*, Western blot analysis of the expressions of apoptotic factors Survivin, Bcl-2 and c-myc. *D-E*, Western blot analysis of the expression of E-cadherin and α-SMA in embryonic pulmonary epithelial cells. The data were measurement data and expressed as mean ± standard deviation. \**p* < 0.05 *vs*. the rats transfected with miR-214 NC and PlGF NC. #*p* < 0.05 vs. the rats transfected with miR-214 NC and PlGF. Data comparisons were performed by repeated measures ANOVA, followed by a Tukey’s post hoc test for multiple comparisons. The experiment was repeated three times. STAT3, signal transducer and activator of transcription 3; PlGF, placental growth factor.

## DISCUSSION

BPD is a respiratory condition occurring in preterm neonates and can lead to chronic respiratory problems driven by several prenatal and/or postnatal factors (19). The known risk factors associated with BPD development in preterm neonates include small gestational age, preeclampsia, chorioamnionitis and infiltration of the chorioamnion by neutrophils (10, 18). The most likely underlying pathogenesis is the constant inflammation in lung, and thus corticosteroid, which has a strong anti-inflammatory effect, has been employed in the treatment of BPD (9). In order to highlight a novel therapeutic method to prevent and cure the pulmonary angiogenesis and alveolarization in neonatal infants with BPD, we planned the current study, and the *in vitro* and *in vivo* results demonstrated that miR-214 could inhibit pulmonary angiogenesis and alveolarization in neonatal infants with BPD *via* PlGF-dependent STAT3 signaling pathway blockade.

Our findings illustrated that miR-214 was downregulated in hyperoxia-induced BPD neonatal rats. MiR-214, a member of microRNA precursors, plays a pivotal role in the pathogenesis of multiple human disorders, including cardiovascular diseases and cancers (26, 38). The expression of miR-214 increased by TWIST1 promotes the epithelial-to-mesenchymal transition and metastasis in lung adenocarcinoma (16). Additionally, our study portrayed that PlGF, which was highly expressed in the lung tissues of preterm rats with BPD, was negatively targeted by miR-214 and activated the STAT3 signaling pathway. PlGF was found to be highly expressed in the BPD rats (33). Moreover, an inverse correlation has been detected in our study between miR-214 and PlGF, and the post-transcriptional miR-214 possesses the ability to modulate the expression of PlGF in lung tissues (12). On the basis of a prior study, PlGF could increase the phosphorylation of STAT3 (2). MiR-214 has been proven to downregulate the expression of STAT3 in human cervical and colorectal cancer cells (4).

Moreover, another critical finding of our study was that miR-214 overexpression could downregulate the expression of IL-1β, TNF-α and IL-6 and repress pulmonary angiogenesis and alveolarization in hyperoxia-induced BPD neonatal rats. IL-1β is one of the main mediators of inflammation and plays a causative role in innumerable diseases (24). In the primary pathological features of BPD, IL-1β contributes to excessive alveolar elastogenesis through the interaction with αvβ6 which serves as an epithelial or a mesenchymal signaling molecule (29). TNF is a kind of ligand related to systemic inflammation of human bodies (7). TNF-α overexpression not only increases the release of glutamate but decreases the cell cycling activity of marrow mesenchymal stem cells (35). IL-6 is a typical proinflammatory cytokine which plays a functional role in a number of physiological inflammatory and immunological processes (1). It has been supported that there is a close correlation between the dysregulation of IL-6 with moderate and severe BPD in preterm infants with a small gestational age (21). PlGF overexpression contributes an exaggerated inflammatory state (17). MiR-214 can inhibit angiogenesis procession by suppressing Quaking and pro-angiogenic growth factor expression (28). Through the depletion of PlGF, Kaempferol exerts suppressive effects on angiogenesis of human retinal endothelial cells (31). Therefore, our studies evidenced that miR-214 overexpression could reduce the expression of pro-inflammatory factors, thus blocked pulmonary angiogenesis and alveolarization as well as inhibit cell apoptosis of lung epithelium in neonatal rats with BPD.

In conclusion, upregulated miR-214 can potentially block the activation of STAT3 pathway by inhibiting its downstream target gene PlGF, ultimately preventing pulmonary angiogenesis and alveolarization in neonatal infants with BPD (Fig. 8). Investigation of miR-214-PlGF-STAT3 pathway in BPD and their functions yields a better understanding of their vivo mechanisms and may have potentially provide important therapeutic implications in the treatment of BPD. However, the molecular mechanism of the miR-214-PlGF-STAT3 pathway in BPD still requires further elucidation with clinical cases involved in the future.

**Fig. 8.**
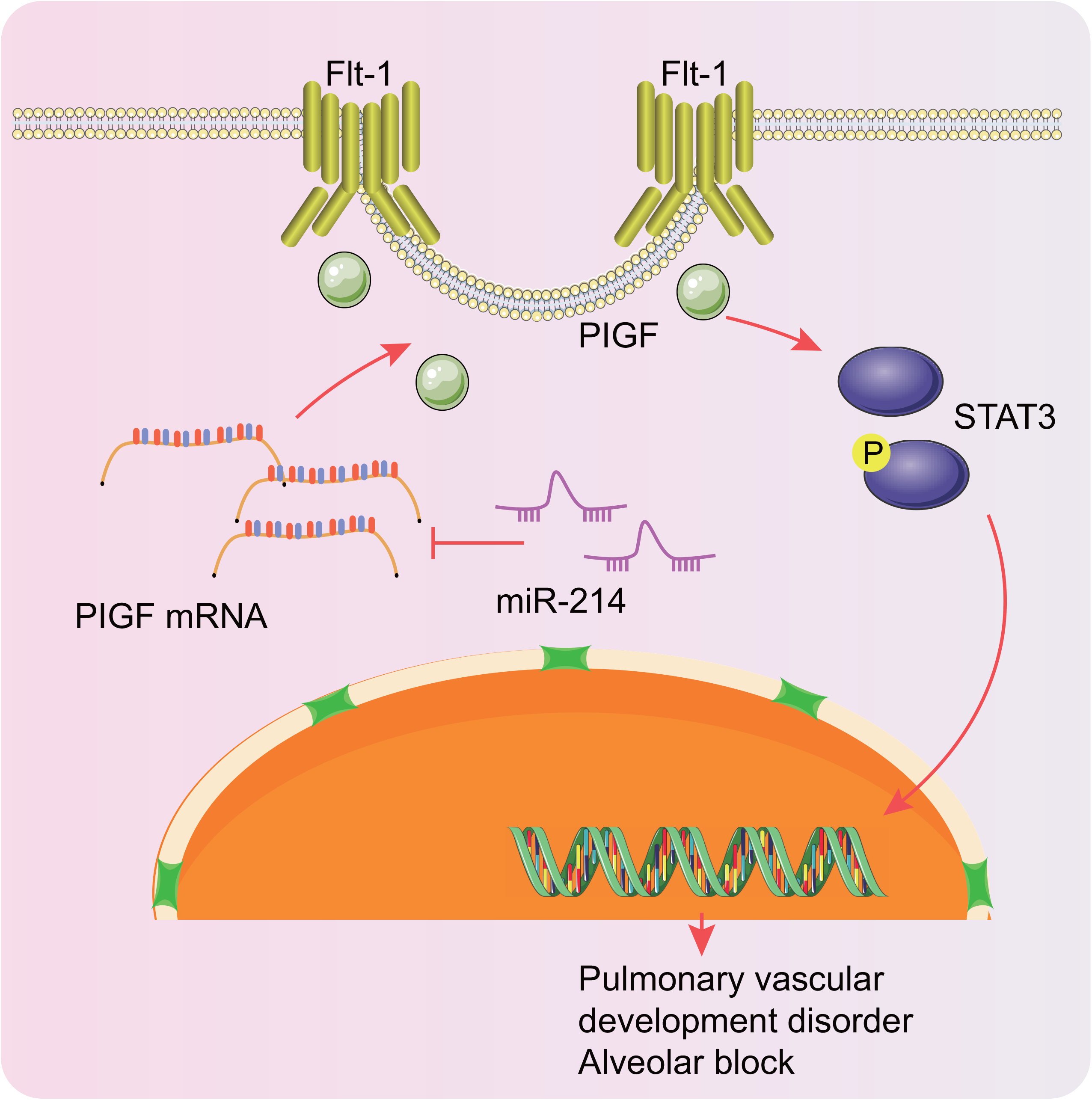
A schematic diagram illustrating the role of the miR-214-PlGF-STAT3 regulatory network in preterm infants with BPD. miR-214 can inhibit the activation of STAT3 signaling pathway by inhibiting the transcription of its downstream target gene PlGF, ultimately impeding pulmonary angiogenesis and alveolarization in preterm infants with BPD. BPD, bronchopulmonary dysplasia; STAT3, signal transducer and activator of transcription 3; PlGF, placental growth factor.

## GRANTS

None.

## DISCLOSURES

The authors have no conflicts of interest, financial or otherwise, to disclose.

## AUTHOR CONTRIBUTIONS

ZQZ, XXL, JL, HH and XMH conceived and designed research; ZQZ and XXL performed experiments; JL and XMH analyzed data; ZQZ and HH interpreted results of experiments; ZQZ and XXL prepared figures; HH and XMH drafted manuscript; ZQZ, XXL and JL edited and revised manuscript; ZQZ, XXL, JL, HH and XMH approved final version of manuscript.

